# Homeostatic Inhibitory Control of Cortical Hyperexcitability in Fragile X Syndrome

**DOI:** 10.1101/459511

**Authors:** C.A. Cea-Del Rio, A. Nunez-Parra, S. Freedman, D. Restrepo, M.M. Huntsman

**Affiliations:** Department of Pharmaceutical Sciences, Skaggs School of Pharmacy and Pharmaceutical Sciences, University of Colorado, Anschutz Medical Campus, Aurora, CO; CIBAP, Escuela de Medicina, Facultad de Ciencias Medicas, Universidad de Santiago de Chile, Santiago, Chile; Department of Cell and Developmental Biology, School of Medicine, University of Colorado, Anschutz Medical Campus, Aurora, CO; Physiology Laboratory, Department of Biology, University of Chile; Department of Pediatrics, School of Medicine, University of Colorado, Anschutz Medical Campus, Aurora, CO

## Abstract

In mouse models of Fragile X Syndrome (FXS), cellular and circuit hyperexcitability are a consequence of altered brain development [reviewed in (Contractor et al., 2015)]. Mechanisms that favor or hinder plasticity of synapses could affect neuronal excitability. This includes inhibitory long term depression (I-LTD) – a heterosynaptic form of plasticity that requires the activation of metabotropic glutamate receptors (mGluRs). Differential circuit maturation leads to shifted time points for critical periods of synaptic plasticity across multiple brain regions (Harlow et al., 2010; He et al., 2014), and disruptions of the development of excitatory and inhibitory synaptic function are also observed both during development and into adulthood (Vislay et al., 2013). However, little is known about how this hyperexcitable environment affects inhibitory synaptic plasticity. Our results demonstrate that the somatosensory cortex of the *Fmr1* KO mouse model of FXS exhibits *increased* GABAergic spontaneous activity, a faulty mGluR-mediated inhibitory input and impaired plasticity processes. We find the overall diminished mGluR activation in the *Fmr1* KO mice leads to both a decreased spontaneous inhibitory postsynaptic input to principal cells and also to a disrupted form of inhibitory long term depression (I-LTD). In cortical synapses, this I-LTD is dependent on mGluR activation and the mobilization endocannabinoids (eCBs). Notably, these data suggest enhanced hyperexcitable phenotypes in FXS may be homeostatically counterbalanced by the inhibitory drive of the network and its altered response to mGluR modulation.

**Significance Statement:** Fragile X Syndrome is a pervasive neurodevelopmental disorder characterized by intellectual disability, autism, epilepsy, anxiety and altered sensory sensitivity. In both *in vitro* and *in vivo* recordings in the somatosensory cortex of the *Fmr1* knockout mouse model of Fragile X Syndrome we show that hyperexcitable network activity contributes to ineffective synaptic plasticity at inhibitory synapses. This increased excitability prevents cortical circuits from adapting to sensory information via ineffective plasticity mechanisms.

## Introduction

Fragile X Mental Retardation Protein (FMRP) is implicated in the transport of approximately 4-8% of all synaptic mRNAs and regulates the translation of numerous proteins involved in synaptic transmission and receptor systems (Brown et al., 2001). Defects underlying FXS are widely believed to lie at the level of the synapse (Zoghbi, 2003; Ebert and Greenberg, 2013) affecting both excitatory and inhibitory neurotransmission across multiple brain regions (Huber et al., 2002; Olmos-Serrano et al., 2010; Paluszkiewicz et al., 2011). Children with FXS exhibit behavioral phenotypes reflective of hyperexcitable circuitry such as a heightened response to somatosensory stimuli (Hagerman and Stafstrom, 2009). This increased/altered somatic sensitivity in FXS is prominent such that, most patients exhibit sensory “defensiveness”, meaning they retreat or pull away when touched (Miller et al., 1999). This hypersensitive phenotype has also been described in the mouse model of FXS [caused by a deletion of the fragile x mental retardation 1 *(Fmr1)* gene] in which *Fmr1* KOs have deficits in whisker-tactile learning tasks due to a hyperactive response to sensory activity (Arnett et al., 2014; He et al., 2017). This altered cortical response may result in poor signal computation, rather than decreased or non-responsiveness to relevant sensory stimulation, supporting the hypothesis that a highly active network fails to adequately decipher sensory inputs in cortex.

The best documented consequence of the absence of FMRP is in inhibition of its function as a repressor to RNA translation subsequent to the activation of mGluRs (Bagni and Greenough, 2005). The “mGluR theory” of FXS [reviewed in (Bear et al., 2004)] accounts for the diverse neurological phenotypes associated with uncontrolled mGluR-mediated protein-synthesis-dependent functions in key brain regions. However, while alterations in excitatory synaptic function and plasticity mediated by mGluRs are well established in the *Fmr1* KO mouse model, little is known about how mGluR-mediated synaptic function and plasticity affecting inhibitory synaptic mechanisms. Electrophysiological and molecular studies identify prominent defects in inhibitory neurotransmission in behaviorally relevant forebrain regions such as the amygdala, hippocampus and cortex of the *Fmr1* KO mice (El Idrissi et al., 2005; D’Hulst et al., 2006; Gibson et al., 2008; Olmos-Serrano et al., 2010; Martin et al., 2014). Considering that sensory input computation and interpretation processes are heavily dependent on local inhibitory network function [reviewed in (Maffei, 2017)], it is important to understand how the aberrant mGluR modulation seen in FXS alters inhibitory circuits of the somatosensory cortex.

In this study, we demonstrate that activation of mGluRs in *Fmr1* KO mice fail to activate inhibitory long term depression (I-LTD). This altered response is primarily due to a decrease in the mGluR-mediated activation of the intracellular mechanisms that lead to this phenomena including endocannabinoid signaling. These data suggest the hyperexcitable phenotype seen in humans is also found in the cortical areas of the mouse brain. Secondly, the *Fmr1* KO mice show an overall disruption of mGluR neuromodulatory and synaptic function, which will have greater impact on specialized interneuron cell populations and inhibitory plasticity phenomena Furthermore, these abnormalities could affect directly the computational processes occurring in layer 2/3 of the somatosensory cortex of the *Fmr1* KO mice disrupting the sensory perception capabilities of the network.

## MATERIALS AND METHODS

### Animals

All experiments were performed under protocols approved by the Ethics and Animal Care Committee of the University of Colorado | Anschutz Medical Campus (Protocol# B-102016(03)1D/00039). Both Control and *Fmr1* KO mice were acquired from the Jackson Laboratories (Bar Harbor, ME, USA) and bred onsite. The *Fmr1* KO animals obtained from these crosses were tested through genotyping protocols. Male mice utilized in this study possess the same congenic FVB background and only hemizygous for the X chromosome *Fmr1* gene were utilized for the experiments in this research.

### Slice preparation

Postnatal 19 to 25 day old control and *Fmr1* KO mice were anesthetized by CO2 inhalation and decapitated. Brains were quickly removed and placed in ice-cold oxygenated sucrose slicing solution composed of (in mM): 234 sucrose, 11 glucose, 26 NaHCO3, 2.5 KCl, 1.25 NaH2PO4 10, MgSO4, and 0.5 CaCl2 (equilibrated with 95% O2 and 5% CO2, pH 7.4). Coronal brain slices (300-μm-thickness) were prepared using a Vibratome (Leica VT1200S, Leica Biosystems, Buffalo Grove, IL, USA). Coronal slices were incubated in pre-warmed (36°C), oxygenated artificial cerebrospinal fluid (ACSF; in mM): 126 NaCl, 26 NaHCO3, 10 glucose, 2.5 KCl, 1.25 NaH2PO4, 2 MgCl2, and 2 CaCl2 for at least 30 min before being transferred to the recording chamber, where they were continuously perfused with ACSF (32°C).

### Electrophysiology

Recordings were obtained in cortical layer 2/3 in control and *Fmr1* KO mice were visually identified using differential interference contrast (DIC) on a modified Olympus upright microscope (Scientifca, East Sussex, United Kingdom). Whole cell recordings were performed with a Multiclamp 700B amplifier (Molecular Devices Corp., Sunnyvale, CA, USA), using recording pipettes (3-5 MΩ) pulled on a PC10 vertical puller (Narishige International, Amityville, NY, USA) and filled with intracellular solution containing the following (in mM): 90 CsCH3SO4, 1 MgCl2, 50 CsCl, 2 MgATP, 0.2 Cs4-BAPTA, 10 HEPES, 0.3 Tris GTP, and 5 QX314 (unless otherwise stated in the manuscript). Recordings were filtered (low-pass) at 4 kHz (Bessel filter) and digitized at 10 kHz (Digidata 1440) using pClamp 10.3 software (Molecular Devices Corp., Sunnyvale, CA, USA). Series resistances were monitored throughout each voltage-clamp recording with 50 ms, −10 mV steps and if it changed by >20%, the data were discarded. Individual inhibitory synaptic events (sIPSC) were recorded in gap free mode in the presence of NMDA and AMPA receptor antagonists (50 μM D-APV and 10 μM DNQX) to block glutamatergic currents and visually identified using pre-written custom code routines in Axograph-X (Berkeley, CA, USA). These events were analyzed by comparing amplitude and frequency between pairing ages in control and *Fmr1* KO mice. For I-LTD and Depolarization-induced suppression of the inhibition (DSI) experiments, evoked IPSCs (eIPSCs) were elicited by 1-ms-long extracellular stimuli using a concentric microelectrode (FHC) placed in layer IV of the somatosensory cortex. Recordings were regularly performed in the continuous presence of NMDA and AMPA receptor antagonists (50 μM D-APV and 10 μM DNQX) unless otherwise stated. Electrical I-LTD was commonly induced after 5 min of stable baseline by high-frequency stimulation (HFS), which consisted of 2 trains (20 s apart), each containing 100 pulses at 100 Hz. On the other hand, chemically induced I-LTD was delivered by application of 10 μM DHPG for 10 minutes while eIPSC were recorded 5 min baseline and 35 min after induction protocol. The magnitude of I-LTD was estimated by comparing averaged responses 35–40 min after HFS with baseline-averaged responses before induction protocol. DSI was evoked by a 1 s voltage step from −60 to 0 mV. eIPSCs were monitored every 4 s for DSI. DSI magnitude was measured as the percentage of change between the mean of the ten consecutive IPSCs preceding the depolarization and the mean of three IPSCs immediately following depolarization (acquired 3–12 s after the pulse).

### In vivo recordings

Tetrode recordings in anesthetized and awake behaving animals were performed as previously described in (Doucette et al., 2011; Gire et al., 2013; Li et al., 2014; Li et al., 2015). Briefly, four tetrodes consisting of four polymide-coated nichrome wires (diameter 12.5 μm, Sandvik) were connected to a 16-channel interface board (EIB-16, Neuralynx) and fed through a housing glued to the board. Immediately before implantation the tetrodes were gold-plated to an impedance of 200-350 MΩ. Adult mice were anesthetized with an intraperitoneal injection of ketamine (100 mg/kg) and xylazine (10 mg/kg). Mice were implanted in layers 2/3 at coordinates AP:-1.46 mm, ML:3 mm. The day of the surgery the optetrode was implanted 200 μm above the final location and every day it was lowered 50 μm until reaching a final depth of DV: 1 mm, respectively. A screw was also implanted in the skull in the opposite hemisphere (1mm right and 2mm posterior of bregma) to serve as ground reference. The animals were allowed to recover at least one week before experiments were performed. The day of the experiment, mice were placed in a 18×12×12 cm anesthesia plexiglass chamber, customized to deliver air puffs trough a port while isofluorane was delivered using a vaporizer. Animal reflex responses were checked constantly through the experiments. Animal were place next to the air puff port and position to get the maximal contralateral neuronal response after whiskers were stimulated with a ~3 L/min air puff. The output of the tetrodes was connected to a 16-channel amplifier (A-M Systems 3500) through a 1× gain headstage (Tucker-Davis Technologies). The signal was amplified 1000× and was recorded digitally at 24 kHz with a Data Translation DT3010 A/D card in a PC computer controlled with a custom MATLAB (Mathworks) program. Spike clustering is explained in detail in (Li et al., 2015). Briefly, data was filtered digitally between 300 to 3,000 Hz. With custom written MATLAB programs, each of the 16 channels was thresholded at three times the standard deviation of the mean. Every spike with amplitude bigger than the threshold was imported into a second program (1 ms record per spike) that performed superparamagnetic clustering and wavelet decomposition of the spikes using 13 different wavelets and three principal components (Quiroga et al., 2004).

## RESULTS

### Decreased neuromodulatory role of mGluRs on inhibitory activity in the somatosensory cortex

As previously reported by us and others (Fanselow et al., 2008; Paluszkiewicz et al., 2011), the mGluR agonist (S)-3,5-Dihydroxyphenylglycine (DHPG, 10μM) increased the frequency of spontaneous inhibitory postsynaptic currents (sIPSCs) in cortical layer 2/3 pyramidal cells. We also found that this increase in frequency was attenuated in *Fmr1* KOs (Paluszkiewicz et al., 2011). To further assess the role of mGluR activity on inhibitory synapses in *Fmr1* KO mice we examined the effect of DHPG sIPSCs in pyramidal cells of layer 2/3 of somatosensory cortex of control and *Fmr1* KO animals. While unnoticed in our previous study where recordings were made at room temperature (Paluszkiewicz et al., 2011), we Initially observe an increase in baseline inhibitory activity as sIPSCs in *Fmr1* KOs were increased (KO: 8.17 +/− 0.46 Hz) in comparison to WT mice (WT: 5.70 +/− 0.55 Hz, p = 0.0011). When DHPG (10μM) was bath applied to pyramidal cells from layer 2/3 of the somatosensory cortex from control animals, sIPSC frequency (from 6.30 ± 0.90 to 11.64 ± 1.20Hz, n = 12, p = 4.5e-5; Fig. 1A and C) and amplitude (from 14.75 ± 1.16 to 21.90 ± 2.47pA, n = 12, p = 0.005; Fig. 1A and D) were significantly increased. These responses were consistently found at DHPG concentrations between 1 to 100 μM, with significant increases as a function of concentration for frequency and amplitude as shown in the dose-response curve in Figs. 1C and 1D. However, when DHPG was bath applied on pyramidal cells from *Fmr1* KO mice there were significantly smaller changes for sIPSC frequency and no changes were observed in amplitude for all concentrations used in the study (at DHPG 10μM: from 7.55 ± 0.55 to 10.16 ± 0.91Hz, p = 0.025, and from 14.08 ± 1.80 to 22.04 ± 4.79pA, p = 0.1; n = 11; Fig. 1B, C and D). When we compared the magnitude of the DHPG effect on control and *Fmr1* KO animals through calculations of the ratio of values for frequency and amplitude after and before DHPG activation, we found that there were significant differences especially for sIPSC frequency in *Fmr1* KO animals (at DHPG 10μM 2.17 ± 0.27, n = 12 vs 1.25 ± 0.19, n = 11; p = 0.03; Fig. 1C), and even for sIPSC amplitude at higher concentrations (at DHPG 100μM: 2.51 ± 0.30, n = 5vs 1.09 ± 0.01, n = 4; p = 0.007; Fig. 1D). Moreover, when the same experiment was analyzed for spontaneous excitatory postsynaptic currents (sEPSCs), DHPG 10μM had similar affects to those observed on sIPSCs, where neither sEPSC amplitude nor frequency was significantly changed in recordings made from pyramidal cells of layer 2/3 of somatosensory cortex of *Fmr1* KO mice (from 10.45 ± 0.48pA to 10.25 ± 0.54pA, n = 5, p = 0.80; and from 15.19 ± 1.48Hz to 15.18 ± 2.01, n = 5, p = 0.99, respectively; Fig. 1E, F and I). Finally, we tested whether other neuromodulators are also functionally disrupted. We bath applied carbachol 10 μM, an agonist of cholinergic receptors, and found that carbachol induced increases of sIPSC frequency (WT: from 4.53 ± 0.54Hz to 7.73 ± 0.18Hz, n = 7, p = 0.0004; KO: from 4.18 ± 0.53Hz to 7.66 ± 0.23Hz, n = 10, p = 0.0001) and amplitude (WT: from 16.67 ± 0.79pA to 52.45 ± 10.28pA, n = 7, p = 0.014; KO: from 16.89 ± 0.92pA to 59.60 ± 10.34pA, n = 10, p = 0.002) from in both control and *Fmr1* KO pyramidal cell recordings (Fig. 1G, H and J). These results suggest that at least the cholinergic neuromodulation on the inhibitory activity of the somatosensory cortex is functional. Altogether, our data indicates that the loss of inhibitory drive activation might be specific for signal transduction associated to mGluRs and not an overall deficiency of the synaptic network.

**Figure 1.**
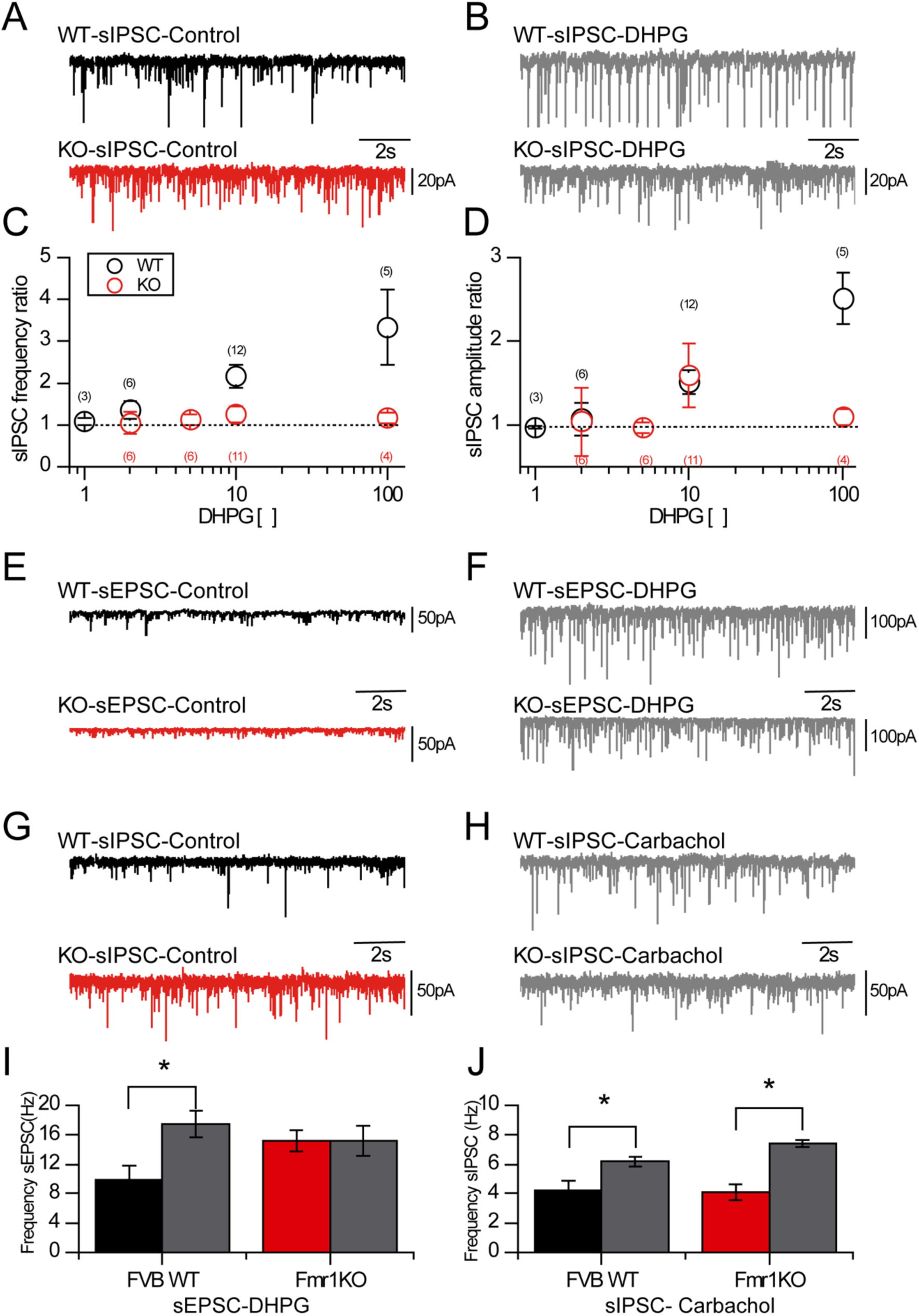
mGluR-mediated activation of the Inhibitory drive in FXS somatosensory cortex. (A) Representative voltage clamp traces for sIPSC activity before application of DHPG 10μM in pyramidal cells from wild type (black) and *Fmr1* KO (red) mouse somatosensory cortex. (B) Representative voltage clamp traces for sIPSC activity after application of DHPG 10μM in pyramidal cells from wild type (upper) and *Fmr1KO* (below) mouse somatosensory cortex. (C) Logarithmic population plot of sIPSC frequency activity ratio after and before application of different concentrations of DHPG recorded from pyramidal cells of wild type (black circles) and Fmr1KO (red circles) somatosensory cortex slices. (D) Logarithmic population plot of sIPSC amplitude activity ratio after and before application of different concentrations of DHPG recorded from pyramidal cells of wild type (black circles) and Fmr1KO (red circles) somatosensory cortex slices. (E) Representative voltage clamp traces for sEPSC activity before application of DHPG 10μM in pyramidal cells from wild type (black) and *Fmr1* KO (red) mouse somatosensory cortex. (F) Representative voltage clamp traces for sEPSC activity after application of DHPG 10μM in pyramidal cells from wild type (upper) and *Fmr1* KO (below) mouse somatosensory cortex. (G) Representative voltage clamp traces for sIPSC activity before application of carbachol 10μM in pyramidal cells from wild type (black) and *Fmr1KO* (red) mouse somatosensory cortex. (H) Representative voltage clamp traces for sIPSC activity after application of carbachol 10μM in pyramidal cells from wild type (upper) and *Fmr1KO* (below) mouse somatosensory cortex. (I) Bar population plot of sEPSC frequency activity recorded from pyramidal cells of wild type (black) and *Fmr1* KO (red) somatosensory cortex slices before application of DHPG 10 μM and after (grey). (J) Bar population plot of sIPSC frequency activity recorded from pyramidal cells of wild type (black) and *Fmr1* KO (red) somatosensory cortex slices before application of carbachol 10 μM and after (grey).

### Heterosynaptic I-LTD is impaired in *Fmr1* KO mice

Several studies have reported the critical roles of mGluR-mediated plasticity in FXS (Nosyreva and Huber, 2006; Bianchi et al., 2009; Zhang et al., 2009; Auerbach and Bear, 2010; Connor et al., 2011; Chevere-Torres et al., 2012; Yau et al., 2016). In order to determine whether the decreased mGluR activation fails to induce synaptic plasticity, we tested the effects of this mGluR malfunction on both electrical and chemically-mediated heterosynaptic I-LTD (Fig.2). I-LTD was measured by analyzing the percentage of change of the integral under the curve of the eIPSC before and after the stimulation protocol. When I-LTD was induced through a protocol of high frequency electrical stimulation (HFS) on layer IV of somatosensory cortex, evoked IPSCs (eIPSCs) from pyramidal cells in layer 2/3 of somatosensory cortex responded with a long-lasting depression (60.14 ± 10.42% of the baseline, n = 8, p = 0.032; Fig. 2A). In contrast, eIPSCs recorded from pyramidal cells of *Fmr1* KO mice did not change in response to our HFS stimulation (88.60 ± 8.04% of the baseline, n = 7, p = 0.29; Fig. 2C), suggesting that I-LTD is impaired in *Fmr1* KOs. To further understand the mechanisms involved in the failure to evoke the electrical-mediated I-LTD in Fmr1KO mice we tested whether mGluRs were critical to this response using a combination of MPEP 4μM (2-Methy-6-(phenylethynyl)pyridine hydrochloride)) and LY367385 100μM ((S)-(+)-α-Amino-4-carboxy-2-methylbenzeneacetic acid) to block all group I mGluRs. When the MPEP/LY cocktail was applied, eIPSCs in pyramidal cells of control mice did not exhibit I-LTD and maintained their strength and amplitude throughout the assay (83.56 ± 5.90% of the baseline, n = 5, p = 0.13; Fig. 2D) These data suggest that group I mGluRs participate in the long-term depression synaptic plasticity process. Moreover, AM251 4μM, an antagonist of cannabinoid receptor (CB-R) also blocked the electrically-mediated I-LTD (96.08 ± 21.01% of the baseline, n = 4, p = 0.92; Fig. 2B), suggesting that the mGluRs are coupled to the activation of CB-R to mediate the depression of the inhibitory drive into pyramidal cells. Furthermore, application of H89 10μM (N-[2-[[3-(4-Bromophenyl)-2-propenyl]amino]ethyl]-5-isoqunolinesulfonamide dihydrochloride), an inhibitor of Protein Kinase A (PKA), during the length of the experiment also blocked I-LTD (97.05 ± 15.06% of the baseline, n = 3, p = 0.83; Fig. 2D), further suggesting that PKA is also involved in the mechanism of this I-LTD response.

**Figure 2.**
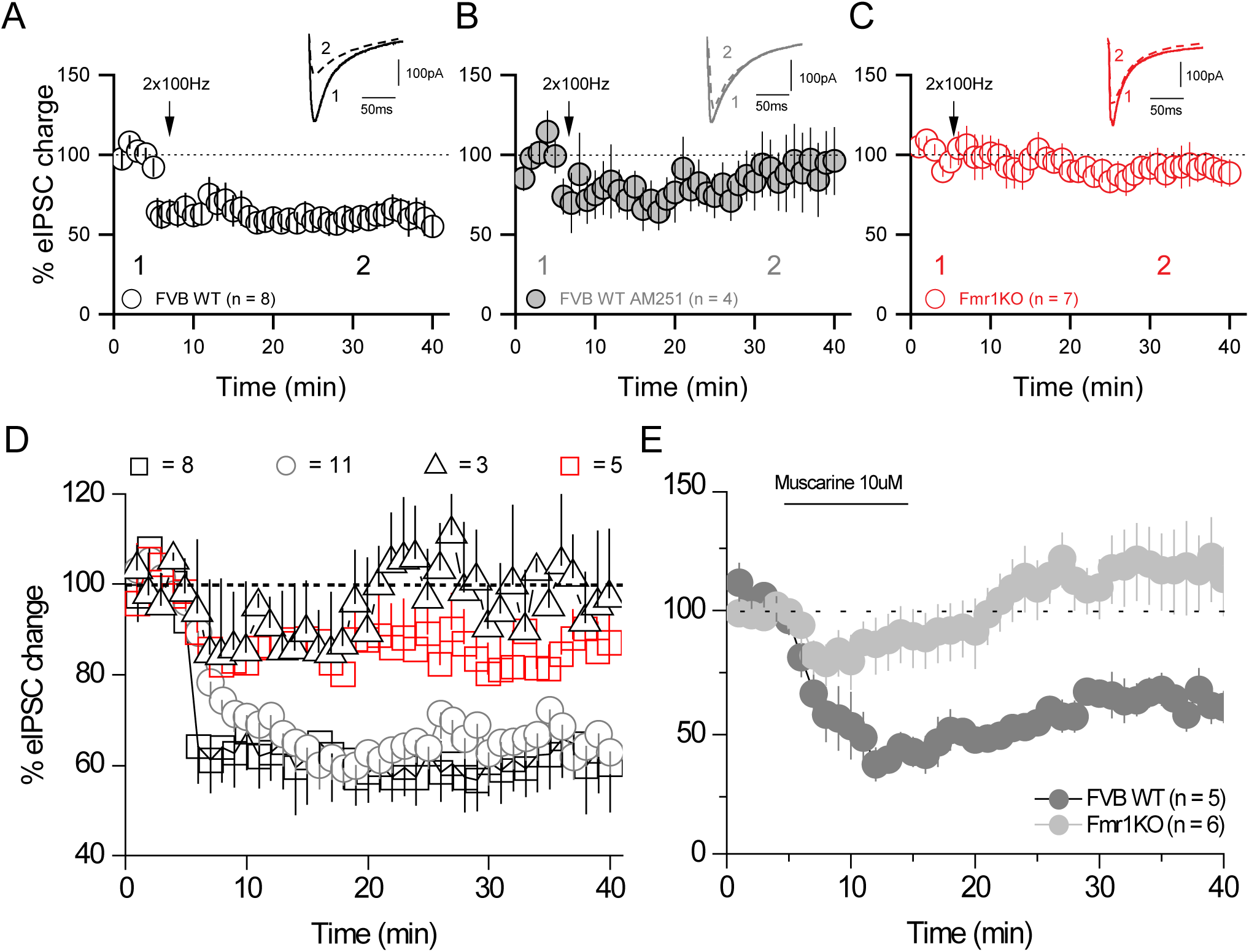
Disrupted electrical-induced heterosynaptic I-LTD in FXS somatosensory cortex. (A) Percentage of eIPSC activity change in the time from pyramidal cell population data recordings in the somatosensory cortex of WT mice before and after an electrical HFS stimulation protocol. Inset shows representative eIPSC waveform before (1) and after (2) HFS protocol. (B) Percentage of eIPSC activity change in the time from pyramidal cell population data recordings in the somatosensory cortex of WT mice before and after an electrical HFS stimulation protocol in the presence of AM251 (an eCB receptor antagonist). Inset shows representative eIPSC waveform before (1) and after (2) HFS protocol. (C) Percentage of eIPSC activity change in the time from pyramidal cell population data recordings in the somatosensory cortex of *Fmr1KO* mice before and after an electrical HFS stimulation protocol. Inset shows representative eIPSC waveform before (1) and after (2) HFS protocol. (D) Percentage of eIPSC activity change in the time from pyramidal cell population data recordings in the somatosensory cortex of WT mice before and after an electrical HFS stimulation protocol (black squares), chemically-induced protocol (grey circles), in the presence of a cocktail of MPEP/LY367385 (red squares) and in the presence of H89, a PKA inhibitor (black triangles). (E) Percentage of eIPSC activity change in the time from pyramidal cell population data recordings in the somatosensory cortex of WT (filled black circles) and *Fmr1* KO (filled grey circles) mice before and after 10 min application of muscarine 10 μM.

Alternatively, we tested whether direct activation of mGluRs could elicit a heterosynaptic I-LTD. Here, we applied DHPG 10μM for 10 minutes while recording eIPSC at 0.25Hz for a total of 40 to 60 minutes. We found that mGluR activation evoked a chemically-induced I-LTD in layer 2/3 pyramidal cells of somatosensory cortex from control animals (62.94 ± 4.71% of the baseline, n = 11, p = 0.0005; Fig. 3A). In order to determine if chemically-induced I-LTD is mediated through endocannabinoid (eCB) mobilization (Chevaleyre and Castillo, 2003), we showed this response is also blocked in the presence of AM251 4μM (99.96 ± 8.60% of the baseline, n = 3, p = 0.61; Fig. 3C) suggesting that eCBs are also mechanistically involved in this type of I-LTD. In contrast, there was a lack of I-LTD in *Fmr1* KO mice (93.60 ± 7.82% of the baseline, n = 7, p = 0.49; Fig. 3B) despite showing significant short term depression (41.81 ± 4.42% of the baseline, n = 7, p = 0.001; Fig. 3B). Altogether these data suggest two important phenomena: 1) that inhibitory drive onto pyramidal cells layer 2/3 of the somatosensory cortex undergoes a heterosynaptic I-LTD in control mice that is similar to previous reports (Chevaleyre and Castillo, 2003), and 2) that I-LTD plasticity is abnormal in the somatosensory cortex of *Fmr1* KOs.

**Figure 3.**
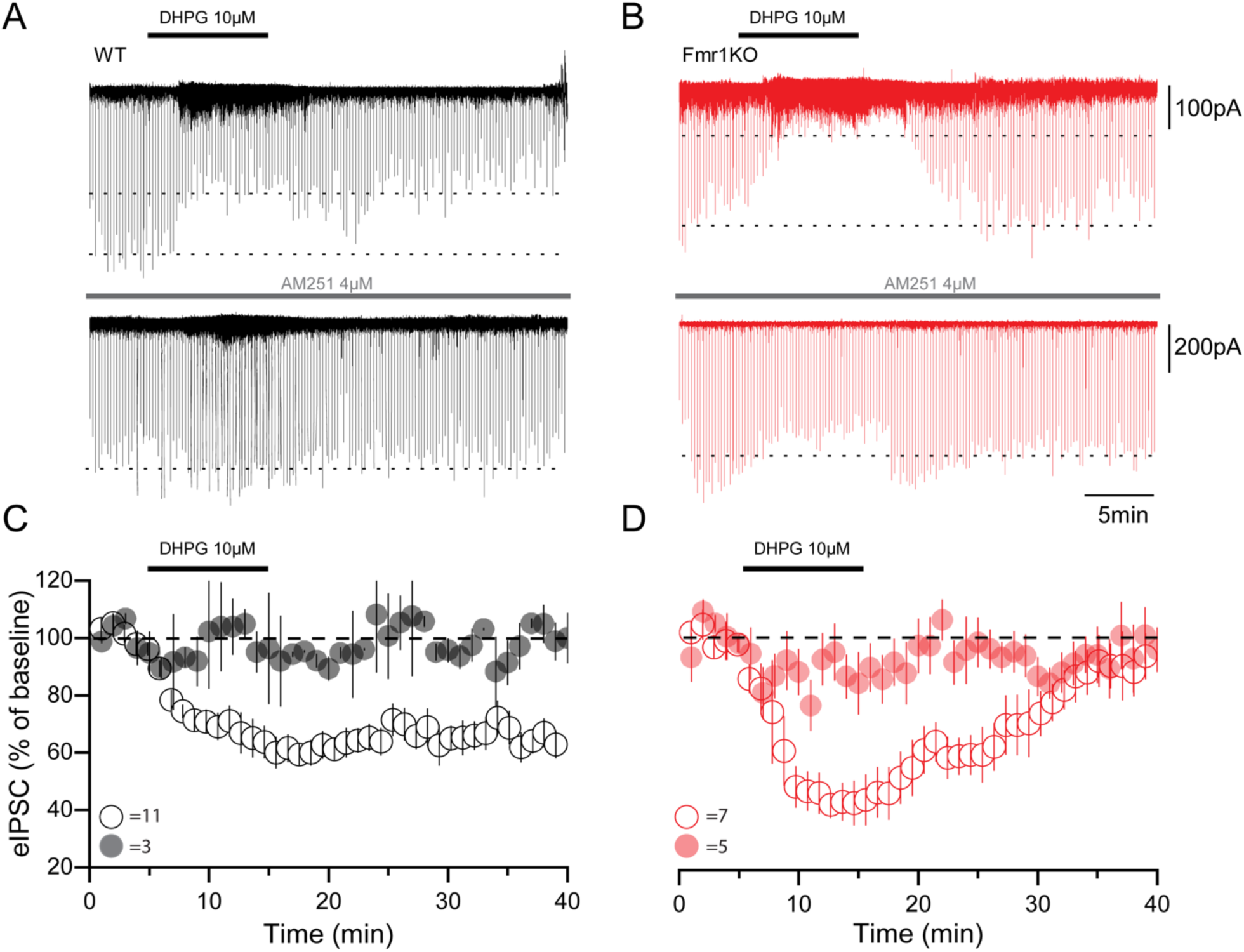
Disrupted chemically-induced I-LTD in FXS somatosensory cortex. (A) Representative traces of DHPG-induced I-LTD in pyramidal cells of WT somatosensory cortex in the absence (upper trace) and presence (lower trace) of AM251 4 μM. (B) Representative traces of DHPG-induced I-LTD in pyramidal cells of *Fmr1KO* somatosensory cortex in the absence (upper trace) and presence (lower trace) of AM251 4 μM. (C) Percentage of eIPSC activity change in the time from pyramidal cell population data recordings in the somatosensory cortex of WT mice in the absence (open black circles) and presence (filled black circles) before and after 10 min application of DHPG 10 μM. (D) Percentage of eIPSC activity change in the time from pyramidal cell population data recordings in the somatosensory cortex of *Fmr1KO* mice in the absence (open red circles) and presence (filled red circles) before and after 10 min application of DHPG.

Faulty I-LTD responses in *Fmr1* KO mice can either be due to lack of direct mGluR activation or failure of the associated molecular signaling pathways. The muscarinic acetylcholine receptor (mAchR) shares common intracellular signaling pathways with mGluRs and they both are able to elicited LTD-I responses (Younts and Castillo, 2013). We tested mAchR activation in pyramidal cells from control and Fmr1 KO mice. Similar to what we found DHPG application in control mice, bath application of the mAchR agonist muscarine (10μM) elicits a long-term depression of inhibitory activity in L 2/3 pyramidal cells (60.79 ± 6.34% of the baseline, n = 5, p = 0.0008; Fig. 2E), thereby supporting the fact that mAchR activation can elicit I-LTD in somatosensory cortex. In contrast, muscarine application onto pyramidal cells of *Fmr1* KO mice failed to elicit I-LTD (113.73 ± 13.98% of the baseline, n = 6, p = 0.51; Fig. 2E). These results suggest that faulty I-LTD is due to a defective intracellular signaling pathway and not solely dependent on the lack of mGluR receptor activation.

### Depolarization induced suppression of Inhibition (DSI) is not altered in Fmr1 KO mice

The I-LTD response is dependent upon endocannabinoid (eCB) mobilization (Fig. 3). To understand other mechanisms of mGLuR-dependent eCB mobilization in *Fmr1* KOs, we examined depolarization suppression of inhibition (DSI). DSI is a known form of short-term plasticity dependent on transynaptic eCB mobilization that is triggered by the depolarization of the postsynaptic neuron resulting in decreased GABA release from the presynaptic inhibitory interneuron (Varma et al., 2001). In these experiments, carbachol (10μM) was bath applied to pyramidal cells to increase the amplitude of the inhibitory drive that is mediated by eCB-sensitive interneurons in the somatosensory cortex. Our results showed no significant differences in DSI. When control pyramidal cells were depolarized from −60 to 0mV for 1 second, sIPSC amplitudes were depressed to a 39.24 ± 7.66% (n = 5) and compared to 51.87 ± 9.38% (n = 9) of the original amplitude in *Fmr1* KOs, indicating no significant differences between KOs and controls (p = 0.32; Fig. 4A, and E). These responses also have similar latencies and time course for control and *Fmr1* KO cells pyramidal cells (Fig. 4D). These results indicate that eCB storage is functional in the *Fmr1* KO mouse model suggesting that the abnormalities seen in I-LTD are not due to storage deficiency of eCBs. Furthermore, when consecutive DSI protocols where applied every 2 minutes, sIPSC amplitude in both control and *Fmr1* KO mice were depressed by the same degree (data not shown), suggesting that either eCB storage capacity is sufficient to elicit DSI or that eCB synthesis rate is not altered in *Fmr1* KO mice. Thus, we conclude that deficiencies in eCBs are not directly responsible in determining the faulty long term I-LTD seen in *Fmr1* KOs.

**Figure 4.**
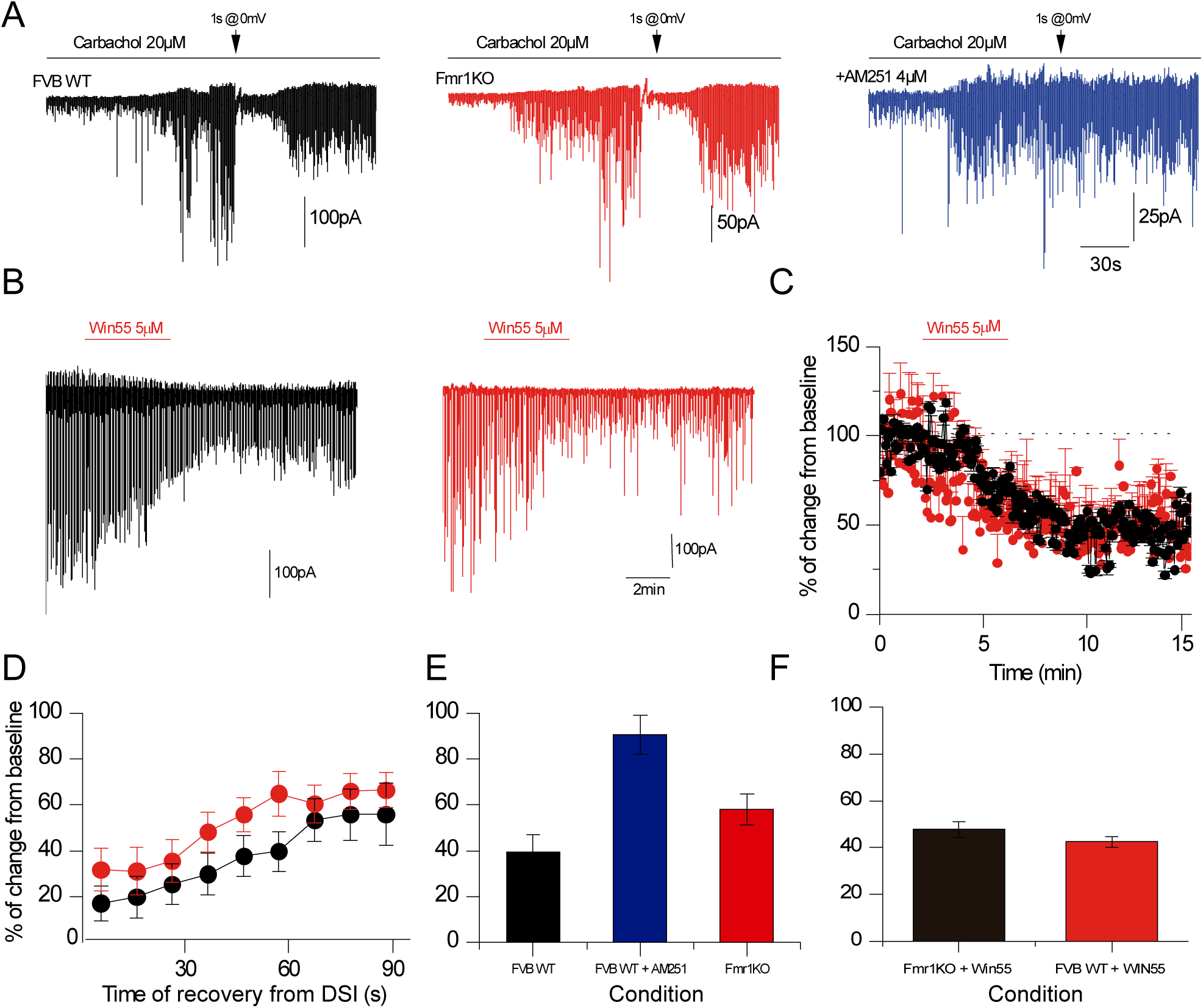
eCB machinery is intact in FXS mice. (A) Representative carbachol induced sIPSC drive before, and after a DSI protocol of 1s of depolarization to 0mV from pyramidal cell recordings of WT (black traces), *Fmr1KO* (red traces) and WT in the presence of AM251 (blue trace) somatosensory cortex slices. (B) Representative traces of eIPSC before and after application of win55-212-2 from pyramidal cells of WT (black traces) and *Fmr1KO* (red traces) somatosensory cortex slices. (C) Data population of percentage of eIPSC change activity from pyramidal cells of WT (black circles) and *Fmr1KO* (red circles) of somatosensory cortex slices in the presence of win55-212-2. (D) Data population of percentage of eIPSC change activity timeline during the next 90s after the DSI induction protocol for WT (black circles) and Fmr1KO (red circles) pyramidal cells of somatosensory cortex. (E) Data population bar plots of maximal percentage of eIPSC change activity after induction of DSI protocol. (F) Data population bar plots of maximal percentage of eIPSC change activity after application of win55-212-2.

Finally, we tested whether eCB receptors were functionally operative in *Fmr1* KO mice testing the effect of Win55-212-2 5μM (an eCB agonist) on eIPSC responses from pyramidal cells. We found that an equal subset of evoked IPSCs (eIPSCs) in control (7/11) and *Fmr1* KO (3/5) pyramidal cells were depressed in response to Win55 (42.47 ± 2.21%, n = 7, p = 1.52e-6, and 54.83± 2.04%, n = 3, p = 0.011), respectively (Fig. 4B and C) suggesting that eCB receptors sensitivity in the *Fmr1* KO is not different to what is seen in control animals.

### Inhibitory response to sensory stimulation recorded *in vivo*

In order to determine how neuronal circuits in layer 2/3 of somatosensory cortex respond to sensory stimulation, we recorded cellular responses in anesthetized and awake behaving control and *Fmr1* KO mice. The setup for tetrode recording in presumed pyramidal cells in layers 2/3 of somatosensory cortex in the anesthetized mouse is shown in Fig. 7A. Off line spike analyses of *in vivo* recordings revealed that in these animals the baseline rate of the recorded units (before stimulation of the whisker) is higher in *Fmr1* KO mice (Fig. 7B, t-test p<0.02, K-S test p<0.005) consistent with hyperexcitability of pyramidal cells in these animals. When the contralateral whiskers are stimulated with an air puff in anesthetized animals, we find that a subset of cells respond differentially (determined with a t-test corrected for multiple comparison of 1 s before the puff and 500 ms after, p<0.05) with either increases or decreases in firing rate (Fig. 7C) in both groups. Importantly, the units in the wild type mice respond differently to those in the in the *Fmr1* KO, with less units exhibiting a decrease in FR after the whiskers were activated (chi-square test, units decreasing FR: 8 out of 24 vs 3 out of 29, p=0.04; units increasing FR 2 out of 24 vs 4 out of 29, p=0.5). Moreover, out of the units that were statistically significantly divergent after the stimulus, exhibit a stronger tendency albeit not significant, to decrease their FR (Fig. 7D, K-S test p=0.1). These data suggest that the pyramidal cells are hyperexcitable and respond differently to contralateral whisker stimulation in *Fmr1* KO mice.

## DISCUSSION

Our results demonstrate that the somatosensory cortex of *Fmr1* KO mice display an abnormal inhibitory drive with *increased* GABAergic spontaneous activity, a faulty mGluR-mediated inhibitory input and impaired plasticity processes, that could in part mediate the hyperexcitability of cortical neurons found *in vivo.* First, there is an overall diminished mGluR activation in the *Fmr1* KO mice that leads to both a decreased spontaneous inhibitory postsynaptic input to principal cells and also to a disrupted form of I-LTD. This I-LTD is dependent on mGluR activation, eCB synthesis and release, and the further activation of presynaptic PKA. These data suggest defective phenotypes in FXS such as hypersensitivity and hyperexcitability may be homeostatically counterbalanced by the inhibitory drive of the network and its altered response to mGluR modulation.

Here, we are the first to demonstrate that, in contrast to excitatory synapses, LTD in inhibitory (I-LTD) synapses is abolished in the *Fmr1* mice. Although these two LTD processes (excitatory and inhibitory LTDs) follow different mechanistic and intracellular pathways of activation, they have at least one critical point in common – mGluR activation. Our results reveal a faulty I-LTD in the somatosensory cortex of *Fmr1* KOs that requires the activation of mGluRs. This I-LTD also depends on eCB release, receptor activation and PKA function as is suggested by the lack of the I-LTD in control animals when mGluR, eCB receptors and PKA activity were blocked. These are all molecular components of the heterosynaptic I-LTD (Chevaleyre and Castillo, 2003; Chevaleyre et al., 2007), suggesting this phenomenon is similar to the one observed in the hippocampus.

Furthermore, although different reports suggest the possibility of direct participation of a faulty enzymatic complex to synthetize eCB in FXS (Maccarrone et al., 2010; Jung et al., 2012), our results indicate that faulty eCB synthesis is unlikely as the main cause of this faulty heterosynaptic I-LTD in FXS. This is because eCB molecular components are downstream to activation of mGluRs, limiting their participation in the I-LTD to functional mGluRs. Additionally, dysfunctional eCB molecular machinery has been proved to be altered mostly in synaptic processes in excitatory synapses without effects on inhibitory connectivity (Maccarrone et al., 2010; Jung et al., 2012). Finally, our experiments on DSI in FXS mice show that eCB activity is unaltered in terms of release and storage availability of the eCB, as well eCB receptor sensitivity. However, this finding does not discard a role for the synthesis of eCBs, but may further support a secondary role for eCB receptor activation in the faulty heterosynaptic I-LTD of the somatosensory cortex.

Our results suggest that different interneuron classes are contributing to abnormal activity of the somatosensory cortex network of *Fmr1* KO mice due to the diminished sensitivity of the mGluRs. Previous reports indicate that the somatostatin positive low threshold spiking cells (Sst-LTS) are especially sensitive to activation via mGluRs (Fanselow et al., 2008; Paluszkiewicz et al., 2011). Based on those findings, we speculate that the lack of response to DHPG of the sIPSC recorded from pyramidal cells in *Fmr1* KO somatosensory cortex could be due to the loss of mGluR activation sensitivity in the Sst-LTS cells. On the other hand, because the mechanism of heterosynaptic I-LTD requires the eCB molecular machinery to be activated as documented by Chevaleyre and others (Chevaleyre et al., 2007), it is highly likely that the interneuron classes participating in this mechanism of plasticity express eCB receptors. The main candidate for this type of response is eCB-1R-expressing perisomatic-targeting basket cells (BCs) that belong to the 5HT-3 expressing group of interneurons of the cortex, and is also characterized by the expression of cholecystokinin (CCK) as is documented in hippocampus and cortex (Foldy et al., 2007; De-May and Ali, 2013). The abnormal function of these interneuron cell types because a faulty mGluR modulation have different implications in the activity of the cortical network. Although these two cell types, Sst-LTS and CCK-BCs, modulate the network in similar fashion having a role in the fine tuning of the information being processed in the cortex, they have different postsynaptic targets (perisomatic vs dendritic) and timing of the response (short vs long-term). Thus, we speculate that while the faulty modulation on the Sst-LTS cells would lead to an immediate but transient truncated inhibitory control over the excitatory activity of the somatosensory cortex, the unregulated I-LTD mediated by CCK-BCs suggest a loss of the long-term control of the inhibitory drive in the FXS animals.

## Homeostatic correction of excitability

The responses described in this manuscript indicate that cortical network inhibition is attempting to counterbalance the hyperexcitability observed in the *Fmr1* KO mouse model of FXS. At baseline levels, inhibitory activity (sIPSCs) is higher in somatosensory cortex of *Fmr1* KOs suggesting there is an attempt to balance the heightened levels of excitatory activity observed in this disorder (Wang et al., 2017). Additionally, in the long term, inhibition is also enhanced in the form of a faulty I-LTD, which allows a persistent inhibitory drive into postsynaptic cells in periods of long lasting high cortical network activity. Interestingly, the fact that the mGluR activation loss of sensitivity in the *Fmr1* KOs could be explained by the already increased inhibitory drive in the disorder, probably at ceiling levels, which would limit the role of mGluRs on activating the network at both baseline and long-term levels. Therefore, this heightened activity likely comes at a cost of a loss of synaptic plasticity. In conclusion, the FXS inhibitory cortical network is homeostatically balancing the high excitatory profile of the disorder, attempting to provide an adequate range of functional neurotransmission modulation for behavioral-related relevant tasks.

**Figure 5.**
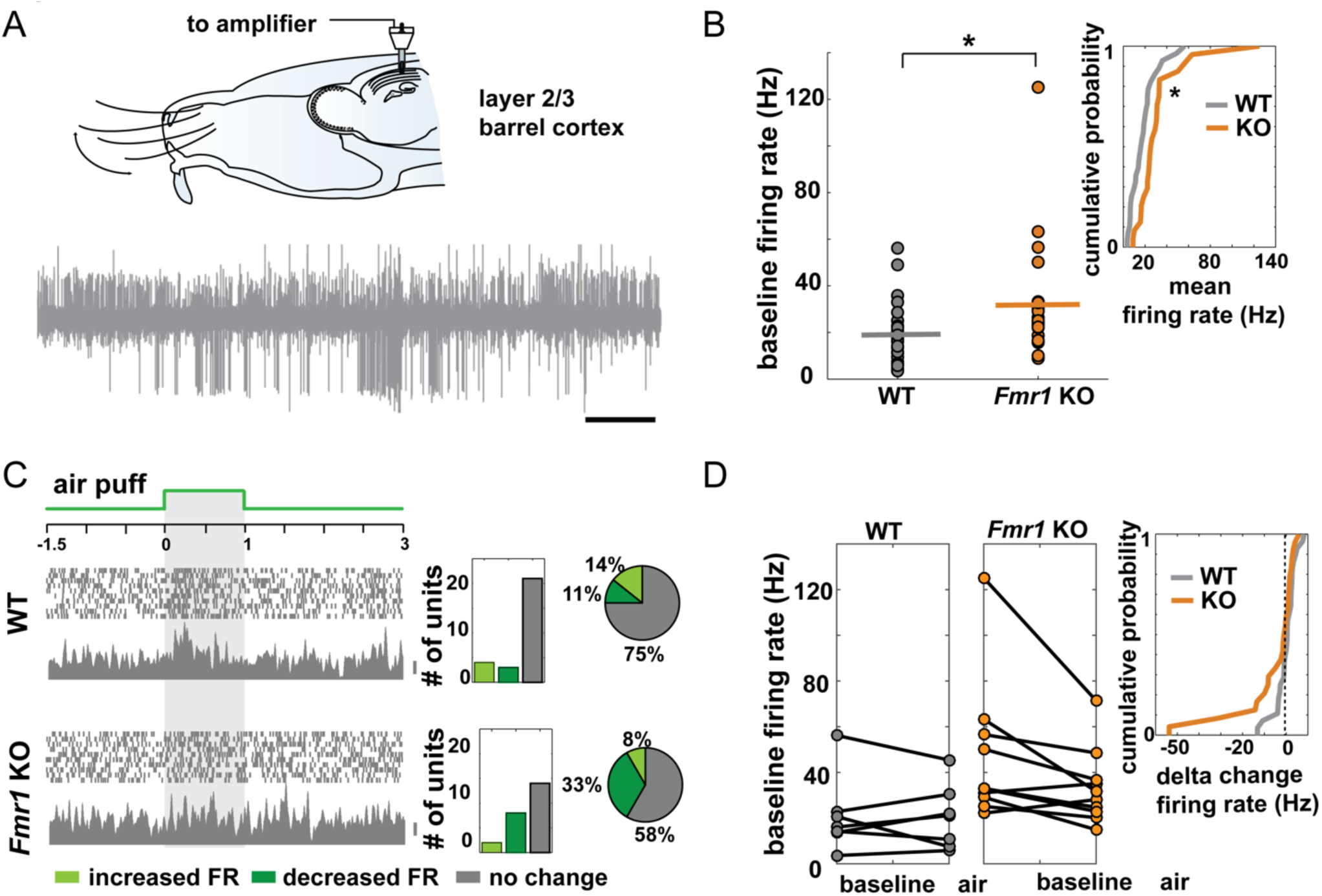
FXS mice cortical neurons of layer 2/3 are hyperexcited *in vivo.* (A) Diagram of the Experimental set-up of in vivo electrophysiological recording. Mice were implants in layer 2/3 with a movable device with 4 tetrodes. Once the animal recovered, there were anesthetized and contralateral whiskers were stimulated with an air puff (~3L/min). (B) Comparison of the basal firing rate of all the neurons recorded in the WT (gray) and *Fmr1KO* (orange). (C) Left, representative excitatory response (10 trials, spikes and PSTH shown, bar 10 spikes) of two units (top panel WT, bottom panel, Fmr1KO) in response to the stimulus. The air puff was delivered for 1 s. Right, population graph depicting the number and percentage of units that exhibited a statistically significant increase, decrease or no change in FR after stimulus delivery. (D) FR before and after stimulus presentation in the WT and KO and cumulative probability analysis of the delta FR (post FR-preFR) in both strains. Right, summary of the neuronal response to air puff determined by t-test (4) (28 units in 2 WT mice and 24 units in 2 *Fmr1* KO mice).

## Acknowledgements

This work was supported by U.S. National Institutes of Health grants (R01 DC000566 to D.R. and R01 NS095311 to M.M.H)

